# Comprehensive transcriptome data of melittin- and un-treated murine hepatoma Hepa 1-6 cells

**DOI:** 10.64898/2026.06.25.734412

**Authors:** Ronghua Zhang, Yuwei Zhang, He Zang, Jiwu Lou, Yueliang Li, Jianrong Jiang, Dafu Chen, Tizhen Yan, Rui Guo

## Abstract

Melittin, the principal bioactive peptide of bee venom, exerts potent antitumor activity against hepatocellular carcinoma (HCC). However, the comprehensive transcriptomic alterations it elicits in hepatoma cells remain poorly characterized. Here, we present an integrated transcriptome dataset from melittin- and un-treated murine Hepa 1-6 hepatoma cells, encompassing messenger RNA (mRNA) and microRNA (miRNA) expression profiles. Cells were exposed to 4 μg/mL melittin in serum-free DMEM for 20 min, and total RNA was subjected to ribosomal RNA-depleted strand-specific RNA sequencing on an Illumina NovaSeq^6000^ platform (paired-end 150 bp) and small RNA sequencing on an Illumina HiSeq^2500^ platform (single-end 50 bp). Raw data were processed using Cutadapt to remove adapters and low-quality reads, yielding clean datasets with *Q*20 ≥ 99.85%, *Q*30 ≥ 98.48%, and valid data ratios exceeding 85%. All raw and processed sequencing data are publicly available. This transcriptomic resource provides a valuable resource and basis for elucidating the regulatory networks underlying melittin-induced anti-hepatoma effects.

**Dataset:** The dataset can be accessed through the National Genomics Data Center, China National Center website by searching with the BioProject accession number PRJCA065485 Reviewers may use this link for anonymous access during the review process. Direct URL to data: Genome Sequence Archive-CNCB-NGDC.

**Dataset License:** CC BY 4.0

## 1. Summary

Hepatocellular carcinoma (HCC) is one of the most prevalent and lethal malignancies worldwide, characterized by high morbidity, mortality, and therapeutic resistance^[1]^. Despite advances in surgical resection, liver transplantation, and molecularly targeted therapies, the prognosis for patients with advanced HCC remains poor owing to tumor heterogeneity, frequent recurrence, and multidrug resistance^[2,3]^. Consequently, novel therapeutic strategies with distinct action mechanisms are urgently needed.

Melittin, a 26-amino-acid amphipathic cationic peptide and the principal active component of western honeybee (*Apis mellifera*) venom, exhibits a broad spectrum of biological activities, including anti-inflammatory, antimicrobial, antiviral, and antitumor effects^[4]^. In the context of cancer, melittin has been demonstrated to induce immunogenic cell death, activate specific T-cell immune responses, modulate macrophage polarization, and trigger mitochondria-dependent apoptosis in various tumor cell lines^[5]^. For instance, in A549 lung adenocarcinoma cells, melittin collapses the mitochondrial membrane potential, triggers a burst of reactive oxygen species, and activates the Bax/Bcl-2-mediated apoptotic pathway^[6]^. Despite these promising findings, the comprehensive transcriptomic landscape and multilayered regulatory mechanisms through which melittin exerts its anti-hepatoma effects remain largely uncharacterized.

To address this gap, we generated a comprehensive transcriptome dataset from the murine hepatoma cell line Hepa 1-6. Log-phase cells were treated with 4 μg/mL melittin (dissolved in serum-free DMEM) for 20 min, alongside untreated controls. Total RNA was extracted using TRIzol reagent, and RNA integrity was verified (RIN ≥ 8.0). For messenger RNA and long non-coding RNA profiling, ribosomal RNA-depleted strand-specific libraries were constructed and sequenced on an Illumina NovaSeq^6000^ platform (2×150 bp paired-end). For small RNA profiling, libraries were prepared with the TruSeq Small RNA Sample Prep Kit and sequenced on an Illumina HiSeq^2500^ platform (single-end 50 bp). All raw data were quality-controlled using Cutadapt to remove adapters, ambiguous bases, and low-quality sequences, achieving high *Q*20/*Q*30 scores and valid data ratios >85%. Clean reads were subsequently used for genome alignment, transcript quantification, and differential expression analysis with standard bioinformatics pipelines. Small RNA reads were also aligned to miRBase v22.1, and unannotated reads were used for novel miRNA prediction.

The final dataset comprises all raw FASTQ files, processed expression matrices, and quality control metrics. All raw data have been deposited in the NCBI SRA under BioProject accession number PRJCA065485. The full raw dataset can be accessed directly via the permanent NGDC BioProject link: https://ngdc.cncb.ac.cn/bioproject/browse/PRJCA065485. No login or access permission is required.

This is the first publicly available comprehensive transcriptome resource profiling melittin treatment in hepatoma cells. It complements existing protein-centric studies by providing a comprehensive map of coding and non-coding RNA alterations, supporting exploration of ceRNA regulatory networks and identification of potential RNA-level therapeutic targets. The high-quality sequencing data can be reused for cross-study comparisons of membrane-active peptides, integrative multi-omics analyses, and development of RNA-directed interventions against HCC.

## 2. Data Description

### 2.1. Dataset Structure

This dataset is deposited in the NGDC (National Genomics Data Center) Genome Sequence Archive under BioProject accession number PRJCA065485. It contains only raw mass spectrometry data files organized as described below. No processed data or additional documentation folders are included in this submission; only the raw data files are provided.

### 2.2 Raw Data Folder Structure

Raw data files are separated by ionization mode and stored in the in the Genome Sequence Archive (Genomics, Proteomics & Bioinformatics 2025) in NGDC for Bioinformation / Beijing Institute of Genomics, Chinese Academy of Sciences (GSA: CRA044082) that are publicly accessible at https://ngdc.cncb.ac.cn/gsa. Each folder contains the following files:

## 3. Methods

### 3.1 Cell culture and melittin treatment

Murine hepatoma Hepa 1-6 cells were acquired from Wuhan Punose Life Science Technology Co., Ltd., and its identity was confirmed through STR profiling to ensure the absence of cross-contamination. The Dulbecco’s Modified Eagle Medium (DMEM), trypsin, PBS, fetal bovine serum (FBS), and penicillin-streptomycin solution were purchased from Gibco, USA. Melittin was acquired from Selleck Chemicals, USA, while RIPA lysis buffer and protease inhibitors were sourced from Shanghai Yamei Biopharmaceutical Technology Co., Ltd. Cells were cultured in high-glucose Dulbecco’s Modified Eagle Medium (DMEM) supplemented with 10% fetal bovine serum (FBS) at 37°C in a 5% CO_2_ incubator. Cells in the logarithmic growth phase were seeded into 6-well plates. For transcriptome profiling, cells were treated with 4 μg/mL melittin (dissolved in serum-free DMEM) for 20 min; control cultures received serum-free DMEM alone for the same duration. For transmission electron microscopy (TEM), cells were similarly treated with 4 μg/mL melittin for 20 min, then harvested and fixed in 2.5% glutaraldehyde at 4°C overnight. After post-fixation with 1% osmium tetroxide, the cells were dehydrated through a graded ethanol series and embedded in Epon resin. Ultrathin sections (70 nm) were stained with uranyl acetate and lead citrate and examined under a TEM (Fig.1).

**Figure 1.**
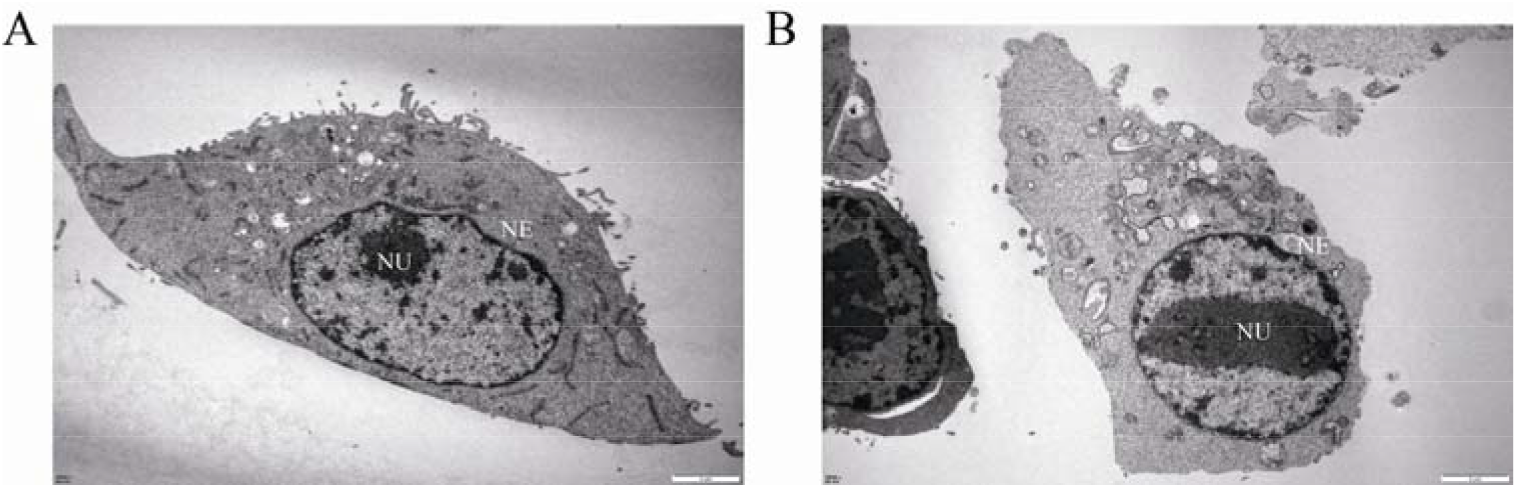
Transmission electron microscopy (TEM) detection of melittin-induced nuclear ultrastructural alterations in Hepa 1-6 cells. (A) Untreated control group, (B) Melittin treatment group at 4 µg/mL (NE: Nuclear Envelope, NU: Nucleolus).

### 3.2 RNA extraction, cDNA library construction, and RNA-seq

By using TRIzol reagent (Invitrogen), total RNA of melittin- and un-treated cell samples were extracted. RNA integrity was detected with an Agilent 2100 Bioanalyzer (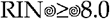). After removing ribosomal RNA, strand-specific libraries were constructed utilizing a dUTP-based method, followed by deep sequencing on an Illumina NovaseqTM^6000^ platform by LC Bio Technology CO.,Ltd (Hangzhou, China).

### 3.3 RNA isolation, cDNA library construction, and sRNA-seq

Small RNA strand-specific libraries were prepared using the TruSeq Small RNA Sample Prep Kits (Illumina, San Diego, USA) and sequenced on an Illumina HiSeq^2500^ platform (single-end 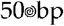) by LC Sciences (Hangzhou, China).

### 3.4 Quality control of raw data generated from RNA-seq

Raw sequencing data were generated using an Illumina sequencing platform, producing 150 bp paired-end reads in FASTQ format. Quality control was performed using Cutadapt (v3.x) to remove adapter sequences and low-quality reads. The following filtering criteria were applied: (i) removal of reads containing adapter contaminants; (ii) exclusion of reads with >10% ambiguous bases (N); and (iii) trimming of reads with quality scores <20 at the 3’ end.

Post-filtering statistics revealed high-quality datasets for both experimental groups. The Control sample yielded 77,800,008 clean reads (11.67 Gb) from 91,042,460 raw reads, with a valid data ratio of 85.45%. The treatment sample produced 79,754,254 clean reads (11.96 Gb) from 90,043,512 raw reads, achieving a valid data ratio of 88.57% (Table 2). Both samples exhibited excellent base calling accuracy, with *Q*20 ≥ 99.85% and *Q*30 ≥ 98.48%, indicating error rates below 0.01 and 0.001, respectively. GC content ranged from 50% to 55.50%, consistent with expected values for *Mus musculus* transcriptomes.

**Table 1.** Contents of the rawdata/raw/neg and rawdata/raw/pos folder.

| File Pattern | Description |
| --- | --- |
| Control raw | Raw data files for control group samples (Control). |
| Control raw.md5 | MD5 checksum files for the corresponding control group .raw files, ensuring data integrity. |
| Treatment raw | Raw data files for melittin-treated group samples (Treatment). |
| Treatment raw.md5 | MD5 checksum files for the corresponding treated group .raw files. |

**Table 2.** Data quality control.

| Sample | Raw Data | Valid Data | Valid Ratio(reads) | Q20% | Q30% | GC content% |
| --- | --- | --- | --- | --- | --- | --- |
| Control | 91042460 | 77800008 | 85.45 | 99.86 | 98.86 | 50 |
| Treatment | 90043512 | 79754254 | 88.57 | 99.85 | 98.48 | 55.50 |

### 3.5 Quality control of raw data produced from sRNA-seq

Raw reads were first checked for integrity using MD5 checksum values, then trimmed using Cutadapt v2.8 to remove adapter sequences, low-quality bases (defined as reads with >20% of bases having a *Q*-score ≤20), poly-A/G tails, and reads containing >5% ambiguous bases (N). The resulting high-quality clean reads were aligned to the miRBase v22.1 database (https://www.mirbase.org/), and unannotated reads were subsequently used for novel miRNA prediction with RNAfold v2.4.12.

Post-filtering statistics confirmed that the quality control procedures yielded high-quality datasets suitable for downstream microRNA expression analysis in both experimental groups. The Control sample generated 13,477,761 valid reads (1.13 Gb) from 22,216,042 raw reads, with a valid data ratio of 60.67%. The Treatment sample produced 8,069,664 valid reads (1.04 Gb) from 20,388,793 raw reads, corresponding to a valid data ratio of 39.58% (Table 3). Base calling accuracy remained excellent throughout the sequencing process: *Q*20 values exceeded 99.37% and *Q*30 values surpassed 97.69%, translating to base error rates below 0.63% and 2.31%, respectively. The GC content ranged from 49.21% to 54.42%, consistent with the expected GC distribution characteristics of *Mus musculus* microRNA transcriptomes. These stringent quality control metrics confirm the reliability of the sequencing data and provide a solid foundation for downstream differential miRNA expression analysis and target gene prediction.

**Table 3.**
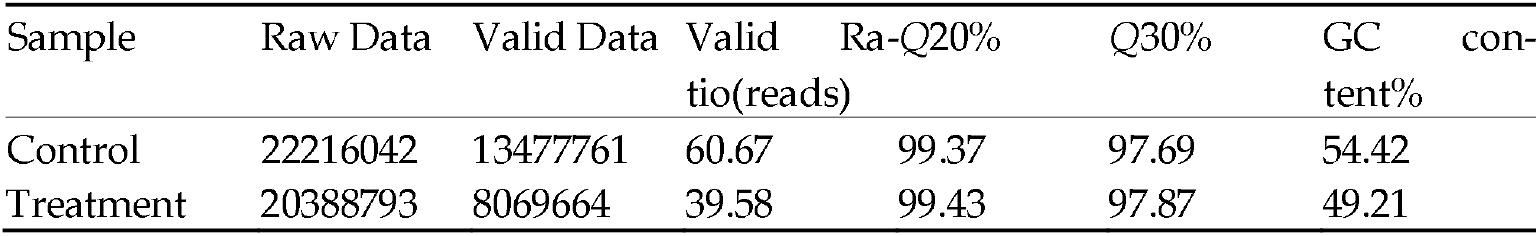
Data quality control.

## Author Contributions

All the authors participated in the conception and design of the study. RH. Zhang, YW. Zhang obtained and analyzed the data. H. Zang, JW. Lou collected the samples. YL. Li, JR. Jiang organized the data and drafted the manuscript. DF. Chen, TZ. Yan, and R. Guo administrated the project, supervised the study, and revised the manuscript.

## Institutional Review Board Statement

Not applicable.

## Informed Consent Statement

Not applicable.

## Data Availability Statement

The raw whole-transcriptome RNA sequencing data, corresponding MD5 checksum files for data integrity validation, and quality control (QC) sample data generated and analyzed in this study are openly available in the National Genomics Data Center (NGDC) Genome Sequence Archive. All raw data are available at the NGDC BioProject PRJCA065485 as described above, under the unique accession number PRJCA065485. No restricted or private data are included in this study.

## Acknowledgments

All of the authors appreciate the constructive and valuable comments from editors and reviewers.

## Conflicts of Interest

The authors declare no conflicts of interest.

The following abbreviations are used in this manuscript:

### Abbreviation Full Name in English

bp: base pair
cDNA: complementary DNA
DMEM: Dulbecco’s Modified Eagle Medium
dUTP: deoxyuridine triphosphate
FBS: fetal bovine serum
FPKM: fragments per kilobase of transcript per million mapped reads
Gb: gigabase
GC: guanine-cytosine
GSA: Genome Sequence Archive
HCC: hepatocellular carcinoma
MD5: Message Digest Algorithm 5
miRNA: microRNA
mRNA: messenger RNA
NCBI: National Center for Biotechnology Information
NGDC: National Genomics Data Center
PBS: phosphate-buffered saline
Q20: Phred quality score 20
Q30: Phred quality score 30
QC: quality control
RIN: RNA integrity number
RIPA: radioimmunoprecipitation assay
RNA-seq: RNA sequencing
SRA: Sequence Read Archive
sRNA-seq: small RNA sequencing
STR: short tandem repeat
TEM: transmission electron microscopy

## Disclaimer/Publisher’s Note

The statements, opinions and data contained in all publications are solely those of the individual author(s) and contributor(s) and not of MDPI and/or the editor(s). MDPI and/or the editor(s) disclaim responsibility for any injury to people or property resulting from any ideas, methods, instructions or products referred to in the content.

## References

1. Wang, S.; et al. Re-exploration of immunotherapy targeting EMT of hepatocellular carcinoma: Starting from the NF-κB pathway. Biomed Pharmacother. 2024, 174: 116566.

2. Huang, H.; et al. RRx-001 inhibits G6PD to deplete NADPH and trigger disulfidptosis coupled with DAMP-mediated immunogenic cell death in hepatocellular carcinoma. Cell Death Discov. 2026, 12, 194.

3. Li, J.; et al. Investigating the role of the TGF-β-SLC20A1 axis in the spatial heterogeneity of hepatocellular carcinoma through single-cell and spatial transcriptomics. Front. Immunol. 2026, 17: 1723334.

4. Król, G.; et al. Functional reprogramming of melittin by Pluronic® F-127 enables anticancer selectivity with attenuated hemolytic activity. Frontiers in Oncology, 2026, 16: 1823622.

5. Duan, X.; et al. Melittin-incorporated nanomedicines for enhanced cancer immunotherapy. Journal of Controlled Release. 2024, 375, 285–299.

6. Li, X.; et al. Melittin kills A549 cells by targeting mitochondria and blocking mitophagy flux. Redox Report: Communications in Free Radical Research. 2023, 28(1): 2284517.

